# Mitochondria-Associated Transcription Precedes Oxidative Phosphorylation Activation During Human Pre-Implantation Embryogenesis

**DOI:** 10.64898/2026.03.25.714226

**Authors:** Andrew J. Piasecki, Melissa Franco, Fausto Capelluto, Konstantin Khrapko, Jonathan L. Tilly, Dori C. Woods

**Affiliations:** Department of Biology, Northeastern University, Boston, MA, United States; Department of Pediatric Oncology and Department of Population Sciences, Dana-Farber Cancer Institute, Boston, MA, United States; Harvard Medical School, Boston, MA, United States; Broad Institute of MIT and Harvard, Cambridge, MA, United States

**Keywords:** Mitochondria, Retrograde Signaling, Embryogenesis, Specification, Differentiation

## Abstract

Mitochondria undergo significant structural and functional changes during human pre-implantation embryogenesis, yet the transcriptional activity of both nuclear-encoded mitochondria-associated genes and mitochondrially transcribed genes across this developmental window remains poorly characterized. While mitochondria are established as the primary energy source for the early embryo, emerging evidence suggests they may also influence lineage specification through epigenetic regulation and metabolite availability. To investigate this, we reanalyzed two publicly available human single-cell RNA sequencing datasets filtered for mitochondria-associated genes using the MitoCarta 3.0 reference database, with separate analyses conducted on the nuclear-encoded and mitochondrially transcribed subsets. The first dataset spanned individual blastomeres from the oocyte through blastocyst stage, and the second compared trophectoderm and inner cell mass cells isolated from blastocysts. Mitochondria-associated gene expression was sufficient to cluster human blastomeres by developmental stage, with morula and blastocyst stage cells forming well-defined clusters. Mitochondrially transcribed genes were found to be the primary drivers of clustering in earlier developmental stages, while nuclear-encoded mitochondria-associated genes drove clustering at the blastocyst stage. A pronounced shift in the expression of both gene sets was identified at the transition from the 4-cell to the 8-cell stage, with 115 unique differentially expressed genes identified across the two stages immediately following this transition, compared to only 5 across the two prior stages. The timing of this transcriptional upregulation, preceding the known onset of oxidative phosphorylation at approximately the 32-cell stage, suggests a mitochondrial role in early embryogenesis beyond energy production. Analysis of trophectoderm and inner cell mass cells showed that mitochondrial gene expression profiles partially distinguished these two lineages, consistent with known differences in mitochondrial activity between them. These findings suggest that both nuclear-encoded and mitochondrially transcribed gene expression is upregulated prior to the first lineage specification event in the human embryo, potentially contributing to epigenetic regulation and cell fate determination through altered metabolite availability. A limitation of this study is its reliance on transcriptomic data alone; future work incorporating functional metabolite measurements will be needed to establish causality. Nonetheless, these data reframe mitochondria as active participants in early human developmental programming rather than passive energy suppliers.

## 1 Introduction

Pre-implantation embryogenesis is an energetically demanding process during which the developing embryo must generate sufficient energy to survive. Mitochondria, which are small and rotund at the oocyte stage, are largely responsible for meeting this energy demand during the initial stages of embryogenesis (Brinster *et al*., 1973, Leese, 2012). However, too much mitochondrial activity is associated with decreased embryo viability, as outlined in the longstanding quiet embryo hypothesis (Leese, 2002). The apparent dissonance between mitochondria serving as the primary source of energy production and the need to limit their activity was reconciled by the concept of functional quietness: an optimal range of mitochondrial output sufficient to sustain development while avoiding cellular stress and excess generation of reactive oxygen species (ROS) (Leese *et al*., 2022). This balancing act demonstrates the importance of proper, well-regulated mitochondrial function within pre-implantation embryos.

Beyond their role in energy production, mitochondria have recently been proposed to influence pre-implantation embryonic development through additional mechanisms. One such mechanism involves mitochondria acting as shuttles for select proteins and RNAs (Cheng *et al*., 2022, Kumar *et al*., 2018). Mitochondria are known to contain nuclear transcription factors that may alter nuclear transcription upon their release, with changes in mitochondrial membrane potential (ΔΨ_M_) and membrane permeability potentially triggering this release in a regulated manner (Beatrice *et al*., 1980). Supporting this, changes in mitochondrial size, structure, and ΔΨ_M_ have been documented in the pre-implantation embryo, most notably during differentiation of the trophectoderm (TE) from the inner cell mass (ICM) (Hayashi *et al*., 2021, Mohr and Trounson, 1982). Separately, mitochondria have been shown to associate with compartments of maternal mRNA in oocytes, theorized to aid in mRNA storage, translation, and degradation to support fertility in mammals (Cheng *et al*., 2022). While the precise function of these mitochondria-associated mRNAs remains under investigation, their existence demonstrates that mitochondria perform roles beyond energy production during this critical developmental window.

Mitochondria also regulate nuclear transcription through their control of intracellular metabolite concentrations. Many metabolites produced or consumed by the Krebs cycle serve as essential cofactors for nuclear transcription factors and epigenetic modulators, directly linking mitochondrial activity to gene expression (Harvey, 2019). A well-characterized example is the dependence of histone acetyltransferases (HATs) and histone deacetylases (HDACs) on acetyl-CoA, a metabolic intermediate whose intracellular concentration is governed by both glycolytic and mitochondrial activity (Harvey, 2019, Moussaieff *et al*., 2015). Experimentally decreasing acetyl-CoA levels in human embryonic stem cells results in loss of pluripotency, implicating mitochondrial control of acetyl-CoA availability in cell fate determination (Moussaieff *et al*., 2015). Similar regulatory relationships exist for other Krebs cycle intermediates including α-ketoglutarate and NAD+, which serve as cofactors for histone and DNA demethylases and sirtuins respectively (Harvey, 2019, Houtkooper *et al*., 2012, Xiao *et al*., 2012). The metabolic state of the embryo is known to shift throughout pre-implantation development, particularly as the TE and ICM lineages differentiate (Leese, 2012), suggesting that mitochondria-driven changes in metabolite availability may contribute to the epigenetic landscape governing lineage specification. Beyond metabolite regulation, mitochondria also influence cellular signaling through ROS production and calcium homeostasis, providing additional routes by which mitochondrial activity may impact transcription during embryogenesis (Duchen, 2000, Hamanaka and Chandel, 2010, Harvey *et al*., 2002, Rizzuto *et al*., 2012).

Despite the growing evidence for expanded mitochondrial roles during pre-implantation embryogenesis, a systematic characterization of the mitochondria-associated transcriptome across the full span of pre-implantation development has not been undertaken. While functional studies indicate that oxidative phosphorylation becomes the primary source of embryonic energy production at approximately the 32-cell stage, whether mitochondria-associated transcription follows the same developmental timeline has not been examined. Understanding the timing of mitochondrial transcriptional activation is important because transcription necessarily precedes the accumulation of functional protein, and any divergence between the transcriptional and functional timelines could point to mitochondrial roles during earlier stages that have not yet been appreciated. Here, we reanalyzed two publicly available human single-cell RNA sequencing datasets by filtering expression data exclusively for mitochondria-associated genes catalogued in MitoCarta 3.0, allowing for a focused interrogation of mitochondrial transcriptional dynamics across developmental stages. This approach was applied to chart how the mitochondria-associated transcriptome evolves from the oocyte through the blastocyst stage, to identify key transitional points in mitochondrial gene expression, and to determine whether mitochondria-associated transcription is sufficient to distinguish both embryonic stage and cell lineage identity within the pre-implantation embryo.

## 2 Materials and Methods

### 2.1 Data Retrieval

The data presented in this manuscript were retrieved from the National Center for Biotechnology Information (NCBI) public data repository. Reads spanning individual blastomeres across pre-implantation embryonic stages were retrieved from GSE36552 (Yan *et al*., 2013) and reads from trophectoderm and inner cell mass cells isolated from blastocysts were retrieved from GSE205171 (Kai *et al*., 2022).

### 2.2 Bioinformatic Prep

All bioinformatic processing was conducted using Galaxy (usegalaxy.org), an open source web-based platform for data-intensive biomedical research. Processing was based on a Galaxy workflow published by (Batut *et al*., 2018), with adjustments made for organism, read length, and strandedness. Raw fastq files were quality checked with FastQC and adaptors were removed using Cutadapt. Reads were mapped to the human reference genome (hg38) using Bowtie2 (Langmead and Salzberg, 2012), and transcript abundance values were generated as transcripts per million (TPM) using StringTie (Pertea *et al*., 2015). For whole cell transcriptome analysis, gene counts were compiled using featureCounts (Liao *et al*., 2013) and differential expression analysis was performed using DESeq2 (Love *et al*., 2014).

### 2.3 Mitochondria-Associated Expression Analysis

Expression values were filtered to include only genes catalogued in MitoCarta 3.0 (Rath *et al*., 2021) and pathway analysis was derived from the PANTHER database (Thomas *et al*., 2003). Principal component analyses, volcano plots, and heatmaps were generated using the packages listed in Table 1. The 4-cell stage was used as the reference for all pairwise comparisons, as it represented the midpoint of the developmental stages present in the dataset and displayed low inter-embryo variation at that stage. Principal component analyses and heatmap visualizations were used as descriptive tools to assess clustering patterns and expression trends across developmental stages.

## 3 Results

### 3.1 Stage-specific embryo clusters can be identified using only mitochondria-associated genes

To assess mitochondrial progression throughout pre-implantation embryogenesis, principal component analysis (PCA) was conducted on blastomeres grouped by developmental stage to determine whether the mitochondria-associated transcriptome alone was sufficient to cluster blastomeres of the same stage. Stages analyzed included oocyte, zygote, 2-cell, 4-cell, 8-cell, morula, and blastocyst. PCA of oocyte through morula stage showed a gradual left-to-right progression in chronological order, with oocyte through 8-cell stage blastomeres clustering primarily on the left (clusters 1, 2, and 3; 40/44 samples, 90.9%) and morula stage blastomeres clustering primarily on the right (clusters 4 and 5; 14/16 samples, 87.5%), suggesting a steady progression of mitochondrial development across these stages (**Figure 1A**). When blastocyst stage cells were included in the same analysis, they did not follow this pattern and scattered across the plot without forming a distinct cluster (**Figure 1C**). This loss of clustering at the blastocyst stage coincides with the onset of lineage specification and differentiation, and may reflect the divergence of mitochondrial profiles as cells commit to distinct fates.

**Figure 1.**
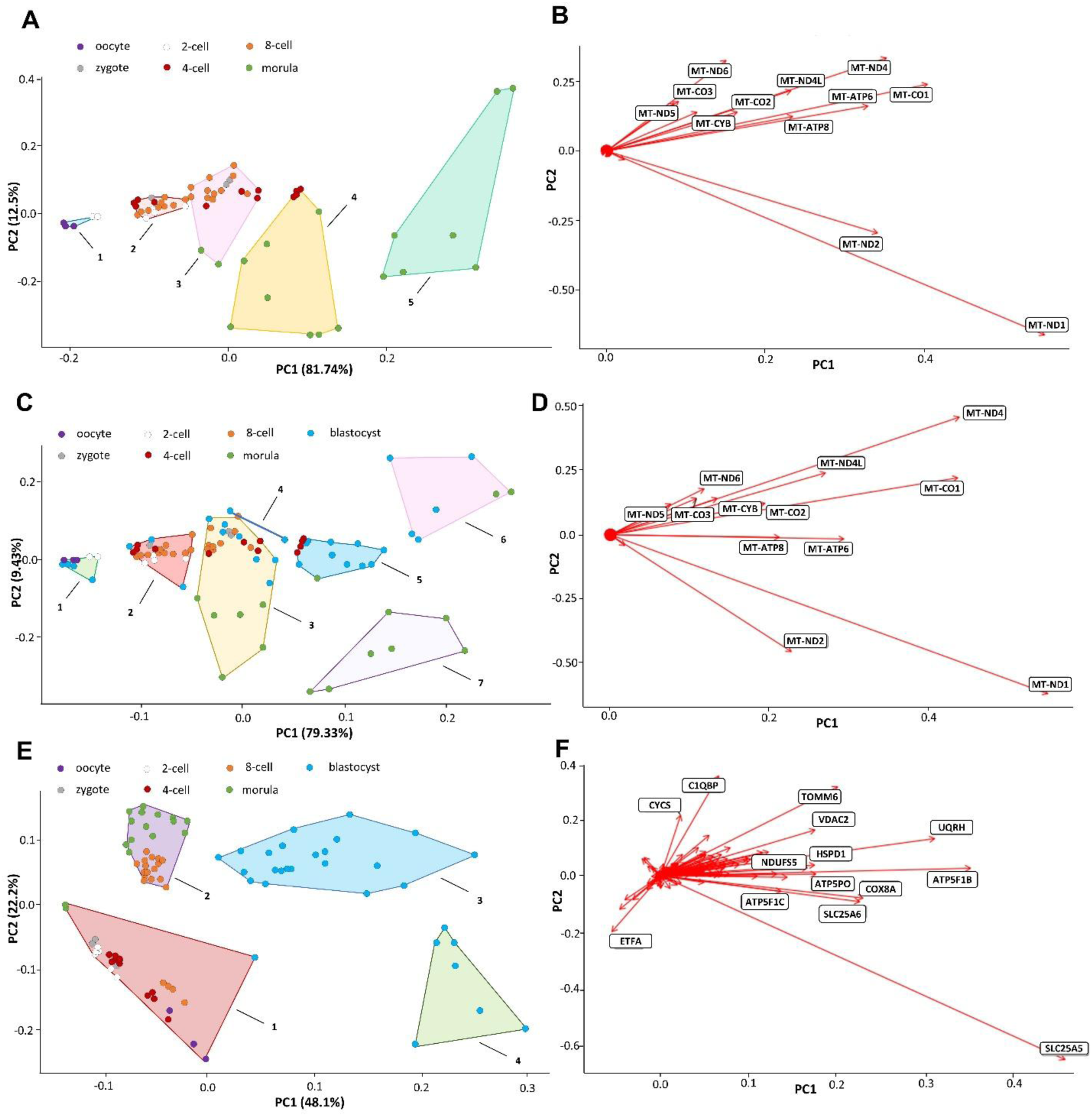
Principal component analysis (PCA) of mitochondria-associated genes at the oocyte (purple), zygote (gray), 2-cell (white), 4-cell (red), 8-cell (orange), morula (green) and blastocyst (blue) stage with the 4-cell stage serving as a baseline. Blastomeres were grouped by embryonic stage. A) Oocyte to morula stage PCA resulted in 5 clusters. Cluster 1 (3/5 oocyte; 2/5 2-cell) was comprised of early stage blastomeres, clusters 2 (1/18 zygote; 4/18 2-cell; 4/18 4-cell; 9/18 8-cell) and 3 (2/19 zygote; 4/19 4-cell, 11/19 8-cell, 2/19 morula) were comprised predominately of blastomeres from the 4-cell and 8-cell stages, cluster 4 (4/12 4-cell; 8/12 morula) was comprised of predominately morula stage blastomeres, and cluster 5 (7/7 morula) was comprised entirely of morula stage blastomeres. B) Primary driver genes for the analysis shown in A, clustering was primarily driven by mitochondrially transcribed genes. C) Oocyte to blastocyst stage PCA resulted in 6 clusters, with the blastocyst stage cells scattering throughout all clusters. Cluster 1 (3/9 oocyte; 2/9 2-cell, 4/9 blastocyst) consisted of early stage blastomeres along with cells from the blastocyst stage, cluster 2 (1/23 zygote; 4/23 2-cell; 4/23 4-cell; 11/23 8-cell; 3/23 blastocyst) was comprised predominately of mid stage blastomeres, cluster 3 (2/28 zygote; 4/28 2-cell; 4/28 4-cell; 9/28 8-cell; 6/28 morula; 7/28 blastocyst) was a transitional cluster comprised cells from multiple different stages, cluster 4 (2/2 blastocyst) is a small cluster of only 2 blastocyst stage cells, cluster 5 (4/15 4-cell; 1/15 morula; 10/15 blastocyst) was predominately comprised of late-stage cells as well as the rightmost mid-stage 4-cell stage blastomeres, cluster 6 (7/7 morula) was a morula cluster, and cluster 7 (2/7 morula; 5/7 blastocyst) was a late-stage cluster. D) Primary driver genes for the analysis shown in C, clustering again was primarily driven by mitochondrially transcribed genes. E) Oocyte to blastocyst PCA with mitochondrially transcribed genes omitted resulted 4 clusters. Cluster 1 (3/31 oocyte; 3/31 zygote; 6/31 2-cell; 12/31 4-cell; 4/31 8-cell; 2/31 morula; 1/31 blastocyst) was the early-stage cluster with a small number of late-stage cells included at the edges, cluster 2 (16/30 8-cell; 14/30 morula) was a mid to late-stage cluster, and clusters 3 (22/22 blastocyst) and cluster 4 (7/7 blastocyst) were blastocyst clusters which notably were very well clustered in this analysis which was not the case in the PCA with mitochondrially transcribed genes included. F) Primary driver genes for the analysis for the analysis shown in E.

In both analyses, mitochondrially transcribed genes were the primary drivers of the clustering patterns (**Figure 1B, D**). To determine whether the same trends held for the nuclear-encoded mitochondria-associated transcriptome alone, PCA was repeated with all mitochondrially transcribed genes removed (**Figure 1E-F**). This resulted in two well-defined blastocyst clusters (29/30 samples, 96.7%), a mixed morula and 8-cell cluster (14/16 morula samples, 87.5%; 16/20 8-cell samples, 80%), and one large cluster containing the remaining 8-cell samples and all earlier stages. Notably, while 87.5% of morula stage samples fell within cluster 2, they accounted for only 46.7% of samples in that cluster, with the remainder being 8-cell stage samples. The presence of 8-cell stage blastomeres within the morula cluster suggests that mitochondrial maturation at the transcriptional level may precede cell division at this stage, with some 8-cell blastomeres already resembling the more mature morula stage profile. These analyses support a significant shift in mitochondria-associated gene expression at the 8-cell stage.

To confirm that the observed differences in mitochondrial profiles between stages were not a result of random inter-sample variation, analysis was repeated with blastomeres from the same embryo grouped independently from those of other embryos at the same stage. Were the variation random, samples of the same stage would be expected to show similar levels of variation between each other as between stages; however, this was not the case. Blastomeres of the same embryonic stage clustered together regardless of embryo of origin and the overall patterns were consistent with the stage-grouped analysis (**Figure 2A-F**), confirming that the stage-specific clustering reflects genuine biological signal rather than dataset bias. As an additional validation, PCA was performed on the full transcriptome. This produced a single cluster containing blastomeres of all stages, with the exception of two morula stage outliers, regardless of whether samples were grouped by stage or by embryo. The absence of stage-specific clustering in the full transcriptome confirms that the patterns identified in the mitochondria-associated transcriptome are not the result of inherent bias within the dataset.

**Figure 2.**
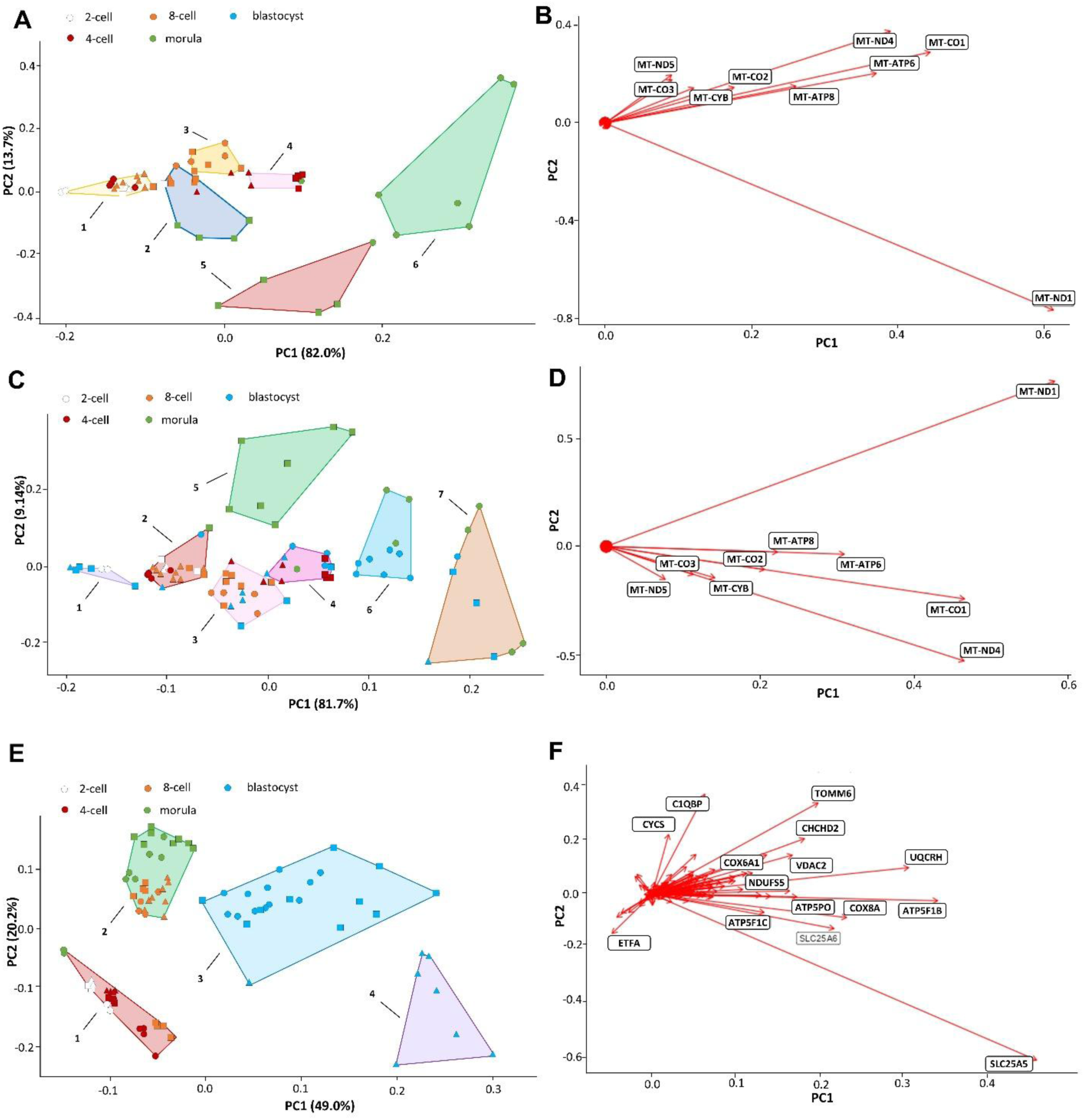
Principal component analysis (PCA) of mitochondria-associated genes at the 2-cell (white), 4-cell (red), 8-cell (orange), morula (green) and blastocyst (blue) stage with a 4-cell stage embryo serving as a baseline. Cells were grouped by embryo of origin. A circle, square, or triangle denotes the embryo of origin for a cell with the stage denoted by the color. A) 2-cell to morula PCA resulted in 6 clusters. Cluster 1 (5/17 2-cell; 4/17 4-cell; 8/17 8-cell) was comprised of multiple early-mid stage blastomeres, cluster 2 (1/11 2-cell; 1/11 4-cell; 5/11 8-cell; 4/11 morula) appeared to be a transitional mid to late-stage cluster, cluster 3 (1/8 4-cell; 7/8 8-cell) was predominately an 8-cell cluster, cluster 4 (6/7 4-cell; 1/7 morula) was predominately a 4-cell cluster, and clusters 5 (5/5 morula) and 6 (6/6 morula) were morula clusters. B) Primary driver genes for the analysis shown in A. As with Figure 1, mitochondrially transcribed genes were the primary drivers of the PCA. C) 2-cell to blastocyst PCA resulted in 7 clusters. Cluster 1 (2/7 2-cell; 5/7 blastocyst) consisted of 2-cell blastomeres and blastocyst cells. It is possible that the blastocyst cells were ICM cells, which would explain low mitochondrial expression similar to that of the earlier embryonic stages. Cluster 2 (4/21 2-cell; 4/21 4-cell; 10/21 8-cell; 1/21 morula; 2/21 blastocyst) contained cells of all stages and is likely a transitional cluster, cluster 3 (2/17 4-cell; 10/17 8-cell; 5/17 blastocyst) also was comprised of various stages, cluster 4 (6/13 4-cell; 1/13 morula; 6/13 blastocyst) contained the 4-cell baseline cells in addition to two other 4-cell blastomeres and some blastocyst cells, cluster 5 (7/7 morula) was a morula cluster, and clusters 6 (3/10 morula; 7/10 blastocyst) and 7 (4/9 morula; 5/9 blastocyst) were morula and blastocyst clusters. D) Primary driver genes for the analysis shown in C, again mitochondrially transcribed genes were the main drivers. E) 2-cell to blastocyst PCA with mitochondrially transcribes genes omitted resulted in 4 clusters. Cluster 1 (6/24 2-cell; 12/24 4-cell; 4/24 8-cell; 2/24 morula) was an early-stage cluster, cluster 2 (16/30 8-cell; 14/30 morula) was a transitional cluster ahead of the blastocyst stage, and clusters 3 (23/23 blastocyst) and 4 (7/7 blastocyst) were blastocyst clusters. F) Primary driver genes for the analysis shown in E.

### 3.2 Development from the 4 to 8-cell stage marks a transitional period in the expression of mitochondria-associated genes

Analysis of mitochondria-associated genes in blastomeres grouped by embryo of origin revealed distinct changes in multiple pathways at key transition points between developmental stages. Volcano plot analysis (**Figure 3A-D**) and heatmap analysis (**Figure 4A-D**) both show an increase in differential gene expression beginning at the 4-cell to 8-cell stage transition. When compared to the 4-cell stage reference, 8-cell stage embryos and morulae showed 75 and 99 differentially expressed genes respectively meeting the criteria of a log2 fold change greater than or equal to 2 and an adjusted p-value less than or equal to 0.01, for a total of 115 unique differentially expressed genes across the two stages immediately following the 4-cell to 8-cell transition. In contrast, the zygote and 2-cell stages showed only 5 and 2 differentially expressed genes respectively meeting those criteria, for a total of only 5 unique differentially expressed genes across the two stages preceding the transition. Pathways showing notable increases in gene expression beginning at the 8-cell stage include oxidative phosphorylation (**Figure 4A**), mitochondrial transcription (**Figure 4B**), and metal and cofactor management. The onset of this transcriptional upregulation at the 8-cell stage, preceding the known activation of oxidative phosphorylation by several cell divisions, suggests that the observed changes in mitochondria-associated gene expression are not solely in preparation for energy production.

**Figure 3.**
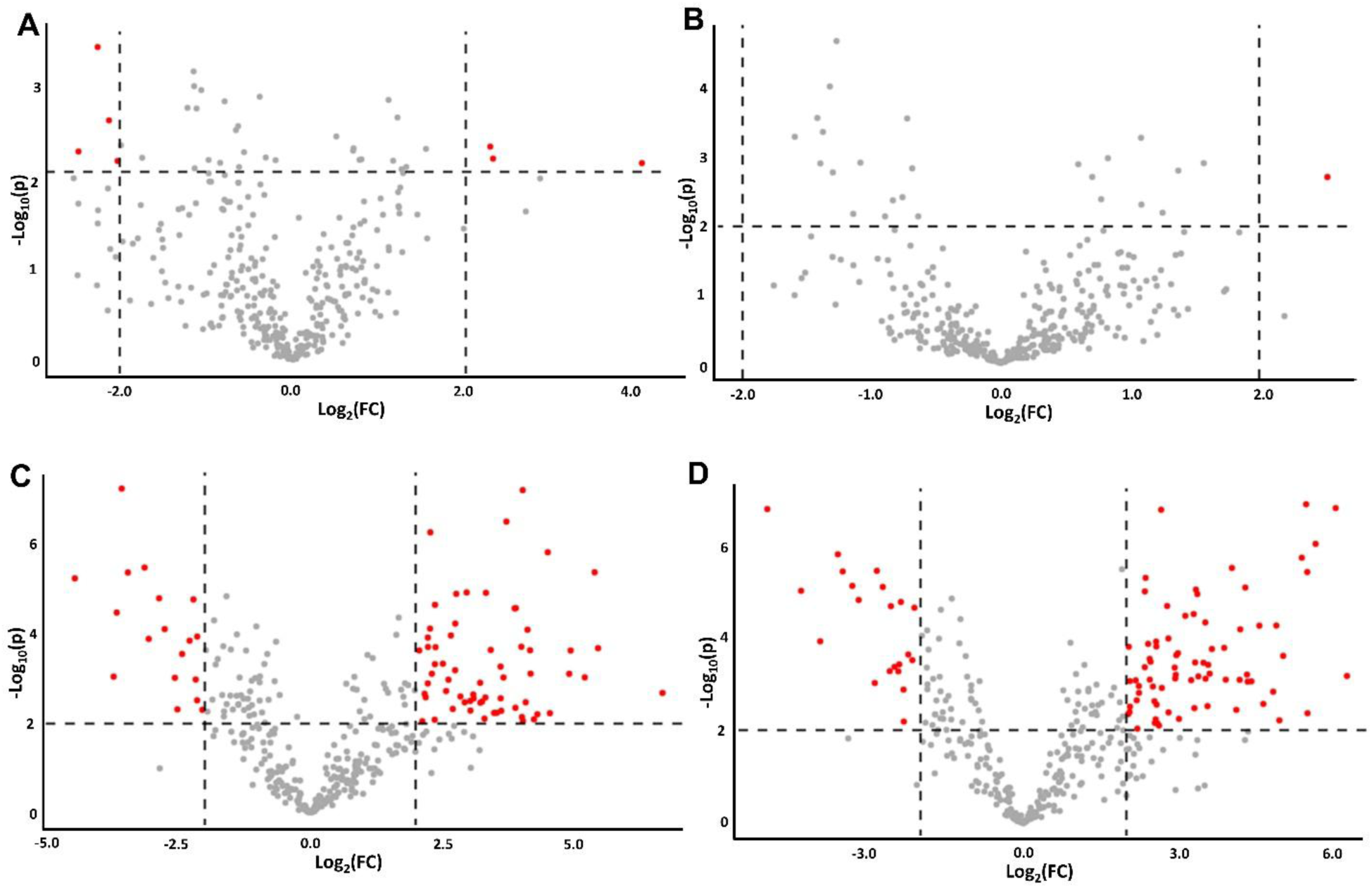
Volcano plots of individual embryos at the A) 2-cell, B) 4-cell, C) 8-cell, D) morula stages using a 4-cell embryo as a reference point. Significant differentially expressed genes were those with a p-value<0.01 and a log2(fold change)>2.

**Figure 4.**
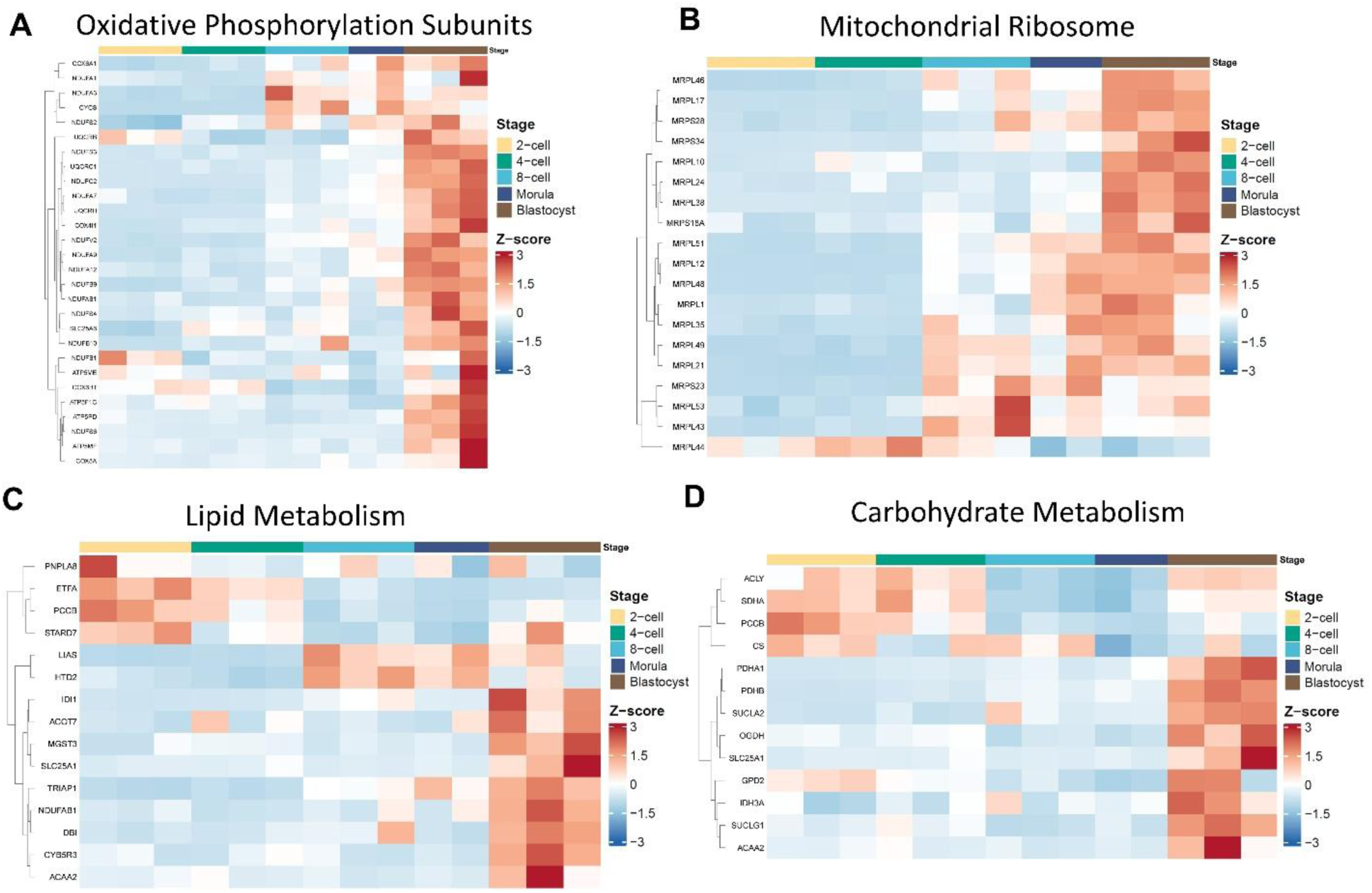
Heatmaps of different mitochondrial-associated pathways shown for each individual embryo. Pathways shown for A) oxidative phosphorylation, B) mitochondrial ribosome, C) lipid metabolism, and D) carbohydrate metabolism. In all cases, shifts in gene expression in these pathways were identified at the 4 to 8-cell transition in addition to the previously well-established morula to blastocyst transition.

Heatmap analysis of carbohydrate and lipid metabolic pathways provides additional evidence that mitochondrial activity shifts in a stage-specific manner throughout pre-implantation embryogenesis (**Figure 4C-D**). Previous studies have demonstrated that metabolite utilization varies across developmental stages (Bradley and Swann, 2019, Haggarty *et al*., 2005, Wale and Gardner, 2012), and the mRNA expression patterns observed here for both carbohydrate and lipid metabolic pathways are consistent with this stage-specific utilization. Importantly, while prior studies have largely characterized preimplantation metabolism through measurement of metabolite consumption rates and energetic output, transcriptomic analysis of the underlying pathway components as presented here provides a complementary approach for identifying the molecular machinery responsible for the metabolic shifts that occur across developmental stages.

### 3.3 TE and ICM cells can be identified using only mitochondria-associated genes

To determine whether the mitochondria-associated transcriptome could distinguish cell lineage identity in addition to developmental stage, a second dataset comprising trophectoderm (TE) and inner cell mass (ICM) cells isolated from human blastocysts (Kai *et al*., 2022) was analyzed using the same MitoCarta 3.0 filtering approach. By the blastocyst stage, cells have committed to either the TE or ICM lineage, and mitochondrial activity is known to differ substantially between these two populations, with TE mitochondria being larger and more active than those of ICM cells, which maintain relatively low mitochondrial activity to limit ROS production (Leese, 2002).

PCA of the filtered expression data showed that mitochondria-associated gene expression partially distinguished the two lineages, with 6 of 9 ICM samples clustering separately from the 9 TE samples (**Figure 5A-F**). While 3 of 9 ICM samples clustered with the TE population, this degree of separation using only mitochondria-associated genes is notable given the relatively small gene set employed and the recency of lineage commitment at this developmental stage. Technical factors inherent to the isolation of ICM from TE under microscopy, including the possibility of residual TE cells adhering to the ICM during dissection, may have contributed to the misclassification of some ICM samples.

**Figure 5.**
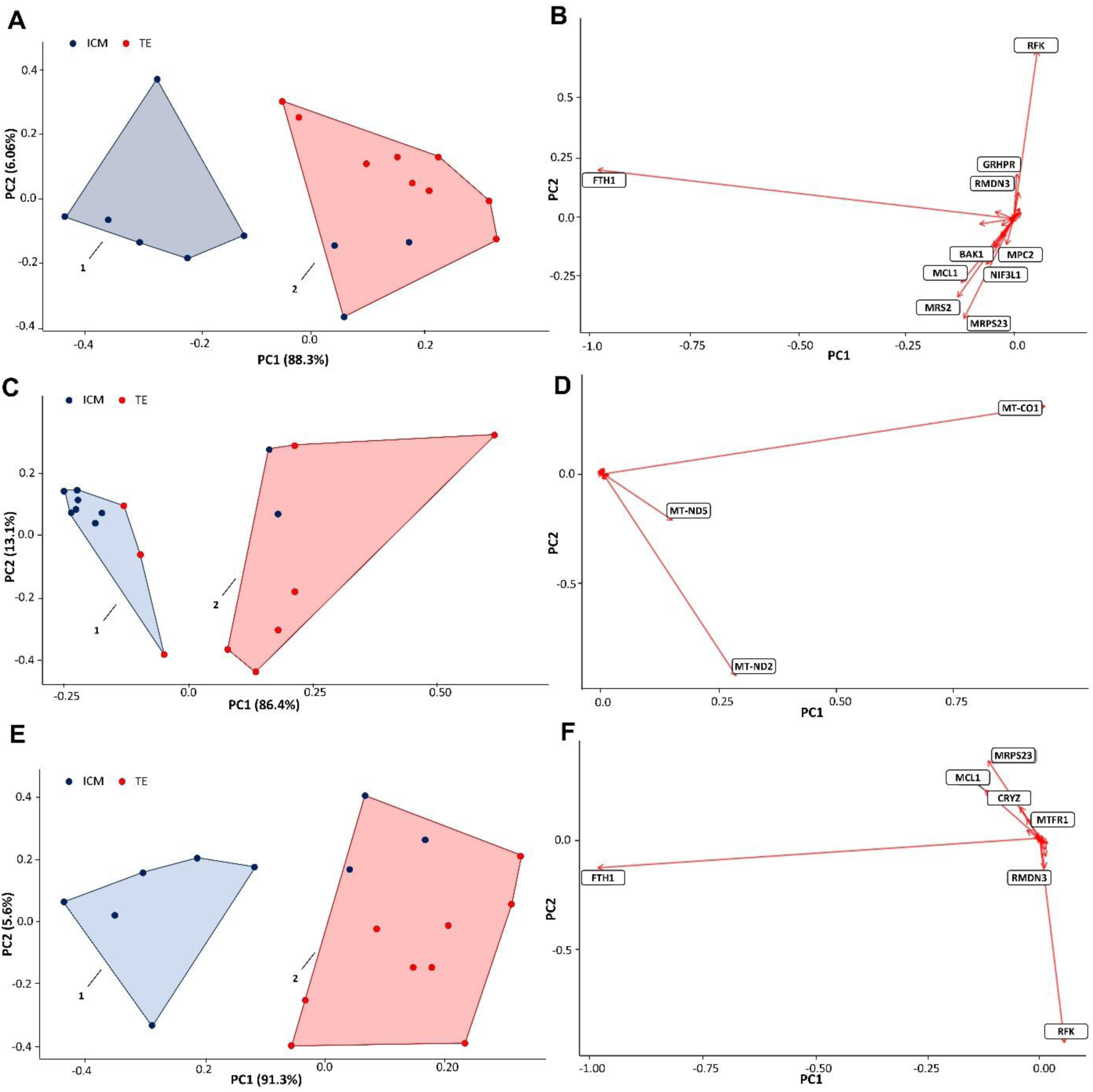
Principal Component Analysis of trophectoderm and inner cell mass samples generated from blastocyst stage embryos. A) ICM cells were set as the baseline and PCA analysis resulted in 2 clusters. Cluster 1 (6/6 ICM) was a pure ICM cluster and cluster 2 (3/12 ICM; 9/12 TE) a predominately TE cluster. B) Primary driver genes for the PCA displayed in A,. Nuclear encoded genes were the main drivers when ICM is set as the baseline. C) TE cells were set as the baseline and PCA analysis resulted in 2 clusters. Cluster 1 (7/10 ICM; 3/10 TE) was the predominately ICM and cluster and cluster 2 (2/8 ICM; 6/8 TE) was the predominately TE cluster. D) Primary driver genes for the PCA displayed in C. Mitochondrially encoded genes were the main drivers when TE is set as baseline. E) TE cells were set as baseline and mitochondrially-encoded genes were removed, which resulted in 2 clusters. Cluster 1 (6/6 ICM) was the ICM cluster while cluster 2 (3/12 ICM; 9/12 TE) was the predominately TE cluster. F) Primary driver genes for TE baseline with mitochondrially-encoded genes removed.

**Figure 6.**
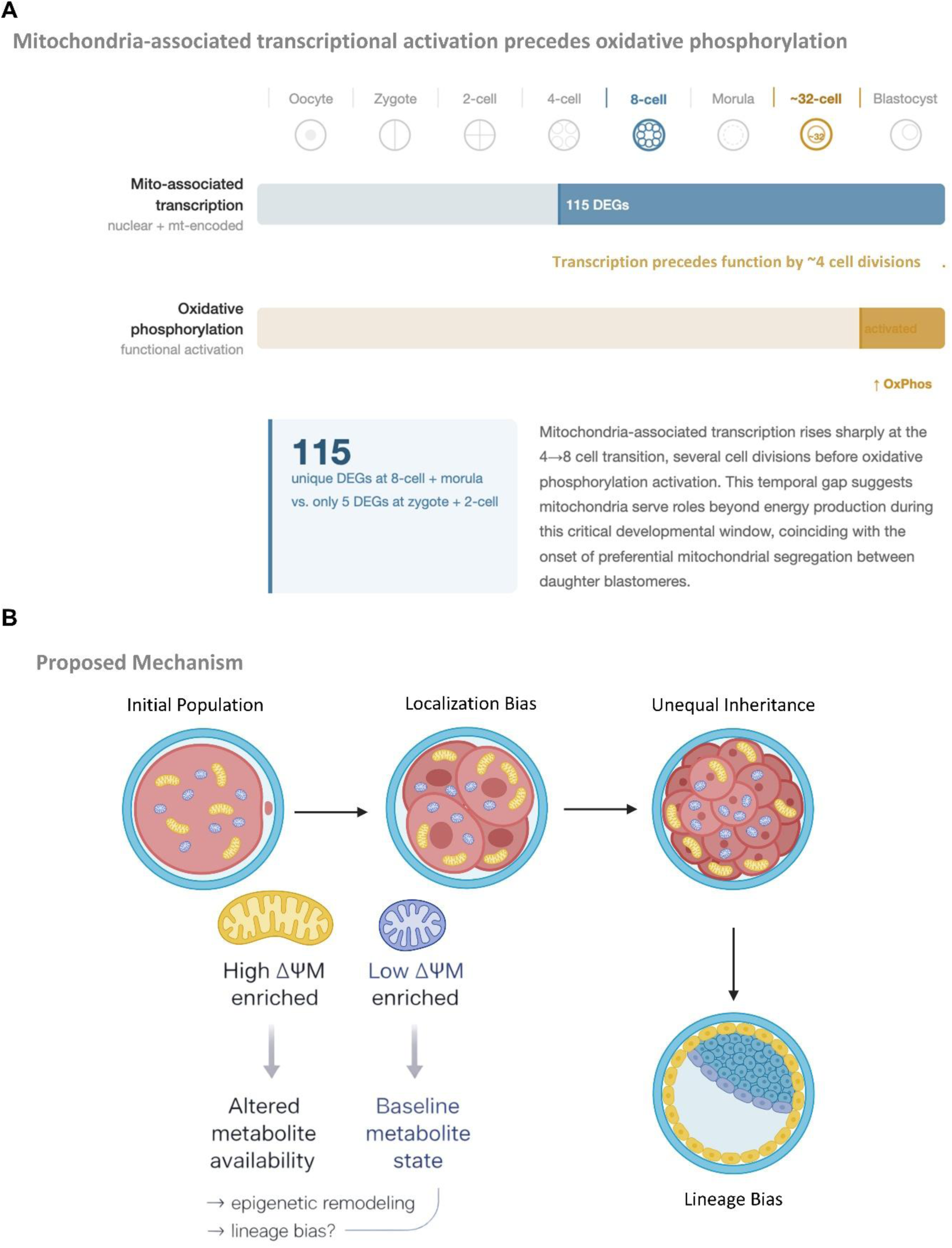
Summary of mitochondria-associated transcriptional dynamics during human pre-implantation embryogenesis and proposed mechanism linking transcriptional remodeling to lineage specification. A) Developmental timeline from the oocyte through the ∼32-cell stage illustrating the temporal relationship between mitochondria-associated transcriptional activation and the onset of oxidative phosphorylation. Transcriptional upregulation of mitochondria-associated genes, evidenced by 115 unique differentially expressed genes identified at the 8-cell and morula stages compared to only 5 unique differentially expressed genes across the zygote and 2-cell stages, is first detected at the 4-cell to 8-cell transition, several cell divisions prior to the known activation of oxidative phosphorylation at approximately the 32-cell stage. Importantly, this transcriptional signal was detectable only through focused analysis of the mitochondria-associated transcriptome; principal component analysis of the full transcriptome produced no stage-specific clustering, indicating that this dynamic would have been masked in aggregate analysis. The temporal gap between transcriptional activation and functional oxidative phosphorylation onset suggests that the observed transcriptional program serves roles beyond preparation for energy production. B) Proposed mechanism by which mitochondrial transcriptional remodeling at the 8-cell stage may contribute to lineage specification. The mitochondrial population of a blastomere is heterogeneous, comprising subpopulations with high mitochondrial membrane potential (ΔΨ_M_, shown in orange) that localize to the outward-facing surface of the cell, and subpopulations with low ΔΨ_M_ (shown in blue) that localize toward the inner contacted surface. Cell division at this stage results in unequal inheritance of these subpopulations between daughter cells. Daughter cells inheriting a greater proportion of high ΔΨ_M_ mitochondria are predicted to have altered metabolite availability with downstream consequences for epigenetic remodeling and lineage bias. For clarity, the schematic depicts a single representative 8-cell stage blastomere. In the intact embryo, mitochondrial heterogeneity and preferential segregation occur across multiple blastomeres simultaneously, with each division having the potential to generate daughter cells with divergent mitochondrial profiles. This panel represents a proposed mechanism supported by existing literature and by the transcriptional findings of this study, and does not reflect a direct experimental finding.

The most informative finding from this analysis was the difference in which gene subset drove clustering depending on the reference lineage used. When ICM cells were set as the reference, clustering was driven primarily by nuclear-encoded mitochondria-associated genes (**Figure 5B**), whereas when TE cells were set as the reference, clustering was driven primarily by mitochondrially transcribed genes (**Figure 5D**). This is consistent with the known biology of these two lineages: the low mitochondrial activity characteristic of ICM cells is reflected in reduced expression of mitochondrially transcribed genes, while the transcriptional differences detectable from the ICM perspective are carried by the nuclear-encoded mitochondria-associated transcriptome. Together these findings suggest that the mitochondria-associated transcriptome retains lineage-specific information even when analyzed in isolation from the broader cellular transcriptome.

## 4 Discussion

Understanding the molecular events governing mitochondrial development during human pre-implantation embryogenesis is fundamental to both developmental biology and reproductive medicine, as this window encompasses the stages at which embryo viability is established and during which embryos are manipulated in assisted reproduction. The findings presented here demonstrate that the mitochondria-associated transcriptome undergoes significant remodeling during this period, with a pronounced transcriptional shift occurring at the 4-cell to 8-cell transition. Previously, the small size and lack of cristae characteristic of mitochondria in early embryonic stages led to the prevailing view that mitochondria were largely dormant until energetically required at the blastocyst stage (Leese, 2002). This view is supported by evidence that mitochondrial DNA (mtDNA) copy number and COX IV (Cytochrome C Oxidase Subunit IV) protein levels do not increase significantly until the blastocyst stage (May-Panloup *et al*., 2005, May-Panloup *et al*., 2007), consistent with the established timeline of oxidative phosphorylation activation. However, the substantial upregulation of mitochondria-associated gene expression observed here at the 8-cell stage, several cell divisions prior to the onset of oxidative phosphorylation, suggests that mitochondrial transcriptional activity is not solely coupled to energy production, and that the functional timeline of mitochondrial activation may be preceded by a transcriptional program that has not previously been appreciated.

Two interpretations of this early transcriptional upregulation warrant consideration. The first is that the embryo begins assembling the molecular machinery necessary for oxidative phosphorylation well in advance of its activation, enabling a rapid upscaling of energy production when energetically required. This interpretation is consistent with the observed increase in oxidative phosphorylation subunit expression at the morula to blastocyst transition (**Figure 4A**), which may represent activation of the final components of the pathway in concert with previously established machinery. The second interpretation is that the transcriptional upregulation at the 8-cell stage reflects mitochondrial functions independent of energy production. The increase in mitochondria-associated transcription at this stage may serve to generate a subset of high membrane potential mitochondria, which are known to be present in the embryo at this stage (Van Blerkom *et al*., 2002) and whose generation cannot be attributed to oxidative phosphorylation activity. Such mitochondria may play roles in transcription factor shuttling or metabolite-driven epigenetic regulation, as discussed in the Introduction. These two interpretations are not mutually exclusive and may reflect parallel processes occurring simultaneously within the developing embryo.

The finding that this transcriptional shift precedes the first lineage specification event is particularly significant when considered alongside the heterogeneous nature of the mitochondrial population within individual blastomeres. The mitochondrial population of a blastomere is not uniform but comprises subpopulations that differ in size, membrane potential, and protein expression (Dumollard *et al*., 2007, MacDonald *et al*., 2019, MacDonald *et al*., 2017). During early embryogenesis, mitochondria with high membrane potential tend to localize to the outward-facing portions of the blastomere, while those with low membrane potential localize toward the inner contacted regions (Van Blerkom *et al*., 2000; Van Blerkom *et al*., 2002; Van Blerkom, 2008).

When a blastomere divides, this spatial organization results in unequal inheritance of mitochondrial subpopulations between daughter cells, with gradients in mitochondrial distribution giving rise to blastomeres that differ in both mitochondrial number and activity (Dumollard *et al*., 2007; Van Blerkom *et al*., 2000). Daughter cells inheriting a greater proportion of high membrane potential mitochondria would have not only increased energetic capacity but also an altered metabolic profile, with downstream consequences for epigenetic regulation and transcription factor availability. The transcriptional upregulation of mitochondria-associated genes identified here at the 8-cell stage — the precise developmental window during which this preferential segregation occurs — provides the first transcriptional evidence that the mitochondrial population is being actively remodeled prior to these fate-determining divisions in the human embryo. Rather than entering lineage-specifying divisions in a transcriptionally static state, blastomeres appear to be actively building a more functionally diverse mitochondrial population, the differential inheritance of which may directly contribute to the divergent developmental trajectories of daughter cells.

The finding that this transcriptional shift precedes the first lineage specification event is particularly significant when considered alongside the heterogeneous nature of the mitochondrial population within individual blastomeres. The mitochondrial population of a blastomere is not uniform but comprises subpopulations that differ in size, membrane potential, and protein expression. During early embryogenesis, mitochondria with high membrane potential tend to localize to the outward-facing portions of the blastomere, while those with low membrane potential localize toward the inner contacted regions (Van Blerkom, 2008, Van Blerkom *et al*., 2000, Van Blerkom *et al*., 2002). When a blastomere divides, this spatial organization results in unequal inheritance of mitochondrial subpopulations between daughter cells, with gradients in mitochondrial distribution giving rise to blastomeres that differ in both mitochondrial number and activity (Dumollard *et al*., 2007, Van Blerkom *et al*., 2000). Daughter cells inheriting a greater proportion of high membrane potential mitochondria would have not only increased energetic capacity but also an altered metabolic profile, with downstream consequences for epigenetic regulation and transcription factor availability. The dramatic upregulation of mitochondria-associated gene expression at the 8-cell and morula stages, immediately preceding the first lineage specification event, is consistent with the hypothesis that mitochondrial heterogeneity contributes to the establishment of distinct cell fates within the pre-implantation embryo.

The mechanisms by which mitochondrial dysregulation disrupts pre-implantation development further support the importance of the transcriptional dynamics identified here. Preimplantation embryos develop in vivo in an environment of approximately 5% oxygen, substantially lower than atmospheric levels of approximately 20%. Exposing embryos to atmospheric oxygen concentrations is sufficient to disrupt mitochondrial pathways and impair embryo viability without reducing energy supply, implicating non-energetic mitochondrial functions including ROS production and metabolite availability as critical determinants of developmental success (Harvey, 2019, Wale and Gardner, 2012). The stage-specific expression patterns identified here for carbohydrate and lipid metabolic pathway components provide a transcriptional correlate for the metabolic shifts that have previously been characterized at the functional level, and may help identify the molecular machinery underlying these transitions.

Taken together, these findings reframe the mitochondria-associated transcriptome as an active and dynamic component of pre-implantation developmental programming, rather than a passive reflection of energetic demand. The identification of the 4-cell to 8-cell transition as a key period of mitochondrial transcriptional activation has potential implications beyond basic developmental biology. The window examined here overlaps directly with the period during which embryos are cultured in vitro during assisted reproduction, and the quiet embryo hypothesis, which has informed embryo selection and culture protocols in IVF, is grounded in the same mitochondrial biology explored here. The stage-specific mitochondrial transcriptional signatures identified in this study may inform future efforts to optimize culture conditions during this critical developmental window and to identify transcriptional markers of mitochondrial health that complement existing morphological assessment approaches. Future work incorporating functional metabolite measurements and protein-level validation will be needed to establish the mechanistic links suggested by these transcriptional data.

## 5 Data Availability Statement

The raw RNA sequencing datasets analyzed in this study are publicly available in the NCBI Gene Expression Omnibus repository under accession numbers GSE36552 (Yan *et al*., 2013) and GSE205171 (Kai *et al*., 2022). Processed differential expression data underlying all figures are provided as supplementary files accompanying this manuscript. The analysis code used to generate figures and conduct mitochondria-associated expression analysis has been deposited to GitHub and is available at https://github.com/ajpiasecki/MPATH. Further inquiries can be directed to the corresponding author.

## 6 Ethics Statement

No new human samples were generated for this study. All data analyzed were retrieved from publicly available repositories. The ethical collection of these samples is described in the original publications from which the datasets were obtained (Kai *et al*., 2022, Yan *et al*., 2013).

## 7 Author Contributions

A.J.P., D.C.W., and J.L.T. conceived and designed the study. A.J.P. performed all data analysis and drafted the manuscript. M.F. wrote the analysis code and contributed to data generation. F.C. contributed to the development of the analytical framework. J.L.T. and D.C.W. interpreted results and critically revised the manuscript. All authors reviewed and approved the final version of the manuscript.

## 9 Acknowledgements

The authors wish to acknowledge Auden Cote-L’ L’heureux for contributions to the finalization of portions of the analysis code used in this study. The underlying analytical framework and pipeline were developed by the authors as part of ongoing work in the Woods laboratory at Northeastern University.

